# FACT mediates cohesin function on chromatin

**DOI:** 10.1101/756601

**Authors:** Jonay Garcia-Luis, Luciana Lazar-Stefanita, Pilar Gutierrez-Escribano, Agnes Thierry, Alicia García, Sara González, Mar Sánchez, Adam Jarmuz, Alex Montoya, Marian Dore, Holger Kramer, Mohammad Mehdi Karimi, Francisco Antequera, Romain Koszul, Luis Aragon

## Abstract

Cohesin is a key regulator of genome architecture with roles in sister chromatid cohesion ^1,2^ and the organisation of higher-order structures during interphase ^3^ and mitosis ^4,5^. The recruitment and mobility of cohesin complexes on DNA is restricted by nucleosomes ^6-8^. Here we show that cohesin role in chromosome organisation requires the histone chaperone FACT. Depletion of FACT in metaphase cells affects cohesin stability on chromatin reducing its accumulation at pericentric regions and binding on chromosome arms. Using Hi-C, we show that cohesin-dependent TAD (Topological Associated Domains)-like structures in G_1_ and metaphase chromosomes are disrupted in the absence of FACT. Surprisingly, sister chromatid cohesion is intact in FACT-depleted cells, although chromosome segregation failure is observed. Our results uncover a role for FACT in genome organisation by facilitating cohesin-dependent compartmentalization of chromosomes into loop domains.

## Main

Sister chromatid cohesion is mediated by cohesin complexes which maintain replicated chromatids in proximity ^9^. Recent studies have shown that cohesins also tether loci in *cis* thus folding individual chromatids into distinct loops that provide an integral level of genome architecture ^3-5^. The current favoured model for how these cohesin-dependent loops are formed on chromosomes involves their dynamic extension through cohesin rings ^3^, also referred to as loop extrusion ^10^. However, direct evidence for cohesin loop extrusion activity has not yet been confirmed. In contrast, single molecule analyses of a related SMC complex, condensin, has shown an ability to form DNA loops in a manner consistent with extrusion ^11^. Cohesin has been shown to translocate along curtains of naked λ-DNA molecules ^7^. Interestingly, the presence of nucleosomes on λ-DNA curtains restricts cohesin mobility ^7^. Since eukaryotic genomes are chromatinised, cohesin functions that involve binding to or translocation along the genome, might require accessory factors to overcome nucleosomes. Recent studies have shown an involvement of the ATP-dependent chromatin remodeller RSC (Remodels the Structure of Chromatin) in the recruitment of cohesion ^6^. Here we set out to identify additional chromatin-remodelers that are important for cohesin function.

We first sought to analyse if nucleosomal occupancy is dynamic at sites of cohesin binding. In *Saccharomyces cerevisiae CEN* sequences are cohesin loading sites where complexes initially access chromosomes before spreading towards pericentric regions 12-15. Genome-wide MNase-seq maps were generated ^16^ in G_1_ and G_2_/M arrests to analyse nucleosome occupancy around centromeric regions (Fig. 1a). Although the yeast genome is characterised by having well-defined nucleosomal positions throughout ^17^, we detected differences in the positioning and occupancy of individual nucleosomes around pericentromeric regions between the G_1_ and G_2_/M MNase-seq maps (Fig. 1a). The general trend involved the loss of defined positioning in G_2_/M arrests (increased nucleosome fuzziness-Fig 1b). The differences were confirmed using DANPOS 2 ^18^ (Fig. 1b) and showed a general correlation with regions enriched in pericentric cohesin (Fig. 1c). Moreover, auxin-mediated degradation of cohesin’s kleisin Mcd1 in G_2_/M arrests shifted the nucleosomal profiles closer to what we had observed for G_1_ arrests (Fig. 1a), i.e. reverting nucleosome fuzziness (Fig. 1b). Together the results are consistent with the idea that nucleosomes are dynamic around *CEN* regions where cohesins are loaded and spread to pericentromeric sites.

**Figure 1.**
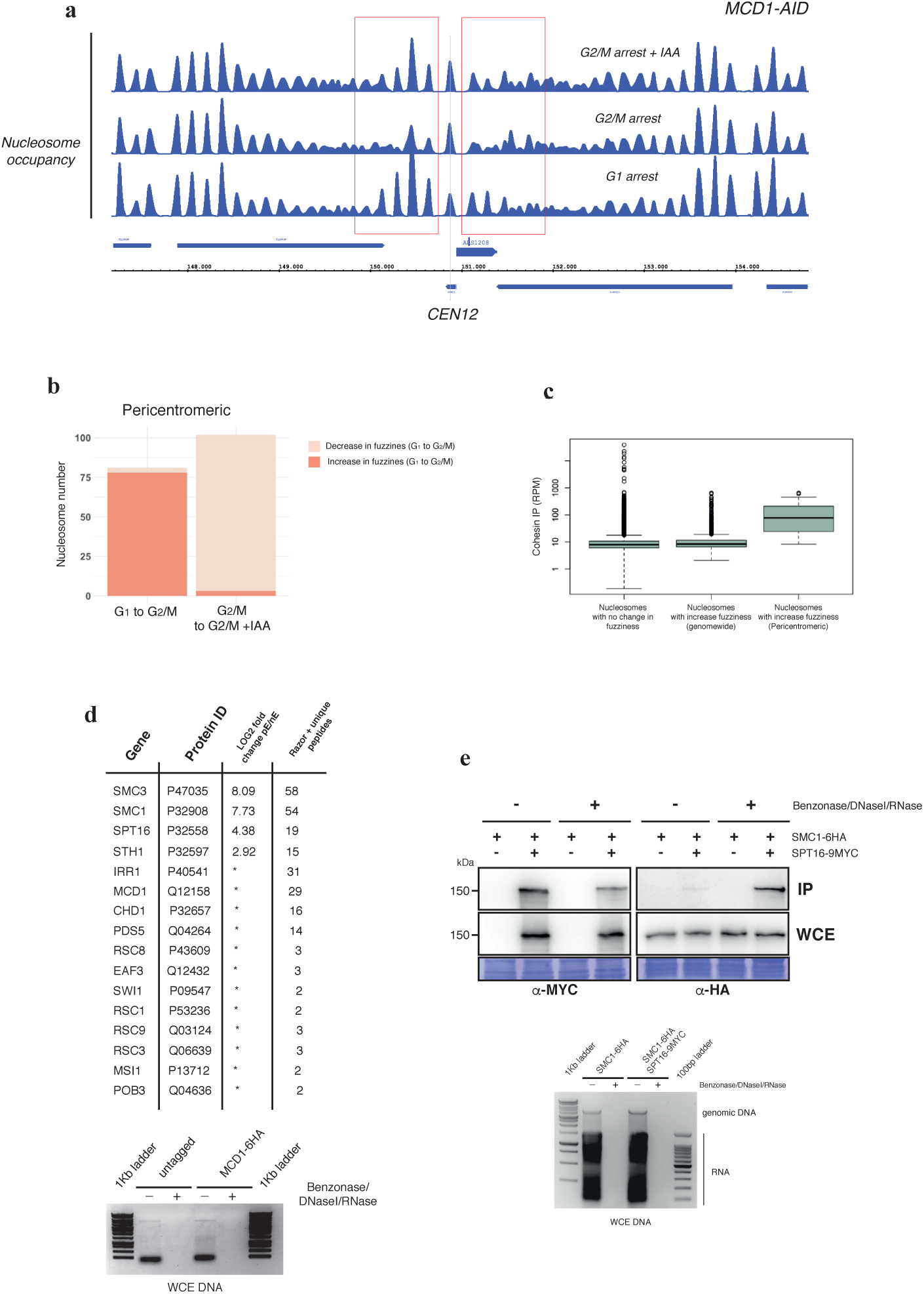
Cohesin complex physically interacts with FACT. **a**, Nucleosome occupancy maps around the centromeric region of chromosome 12 of a strain carrying an auxin degradable allele of Mcd1 (*MCD1-AID*) arrested in G_1_ (G_1_ arrest), in metaphase by nocodazole (G_2_/M arrest), or after auxin-mediated degradation of Mcd1 following G_2_/M arrest (G_2_/M arrest + IAA). Regions with changes in nucleosome occupancy are boxed in red. **b**, Whole genome DANPOS analysis of changes in nucleosome fuzziness between G1 and G2/M arrests. Increased fuzziness indicates number of nucleosomes well positioned in G1 that become fuzzy in G2/M, while decrease fuzziness indicates the opposite. Y axis represent log-scale cohesin enrichment in RPM. Box plot of centromeric nucleosomes is focus on pericentric regions (+/- 10kbp of CEN sequences). Centre lines depict the medians; box limits indicate 25th and 75th percentile. The whiskers will extend up to 1.5 times the interquartile range from the top (bottom) of the box. The data points beyond that distance are represented as ‘outliers’. The differences between the top and bottom whiskers are based on the fact that log transformation of the data does not maintain the distance of a point from the third or first quartile. **c**, Box plot analysis of cohesin enrichment with respect to nucleosome fuzziness in the yeast genome or at pericentromeric regions. Y-axis represents cohesin enrichment in RPM. **d**, Proteomic screen was conducted to identify chromatin remodelers that interact with the cohesin complex in budding yeast. Protein extracts from *GAL-CDC20 MCD1-6HA* and untagged control strains arrested in metaphase were treated with a mix of Benzonase, DnaseI and RnaseI and immunoprecipitated using anti-HA conjugated magnetic beads. IP elutions were subjected to trypsin digestion and analysed by LC-MS/MS. For each protein identified, label-free quantification intensities were compared between *MCD1-6HA* (pE) and untagged (nE) samples and the enrichment was calculated as the log_2_ fold change. The table on the left panel summarises cohesin subunits and chromatin remodelers identified in the analysis, including the enrichment values and the number of peptides found (MQ Razor+Unique peptides). Asterisks in the enrichment column represent proteins identified in the tagged but not in the untagged elution. Bottom panel shows the result of the Benzonase/DnaseI/RnaseI treatment for the degradation of nucleic acids in the protein lysates. **e**, Analysis of cohesin and FACT interaction. Protein extracts were treated with DNAseI/Benzonase/RNAse to remove DNA (bottom) before immunoprecipitation of Spt16-9myc through MYC epitope. Smc1-6HA was detected in the pulldowns, confirming the interaction, in the presence and absence of DNAseI/Benzonase/RNAse treatment.

Next, we sought to identify potential chromatin remodellers involved in cohesin function. We immunoprecipitated Mcd1 from G_2_/M arrests and analysed cohesin interactions by mass spectrometry (Fig. 1d). Among the list of proteins identified we found subunits of several chromatin remodeller complexes. The most abundant was the Spt16 subunit of the FACT complex (<u>fa</u>cilitates <u>c</u>hromatin transcription) (Fig. 1d-e). Consistent with previous reports ^19^, several subunits of RSC were also amongst the hits (Fig. 1d).

FACT has a dual activity destabilising and reconstituting nucleosomes ^20-22^, which facilitates translocation of protein complexes on DNA during transcription and replication ^23^. Importantly, FACT is required for the *in vitro* assembly of mitotic chromosomes by condensin ^24^. These functions make FACT an ideal candidate to facilitate cohesin functions on chromatin. To further explore this, we first sought to test whether inactivation of FACT in G_2_/M arrests caused any alterations in cohesin binding. An auxin (IAA)-inducible degron of Spt16 (*SPT16-AID*) ^25^ was used to deplete FACT from cells in metaphase (Suppl. Fig. 1). We performed calibrated ChIP-seq as described in the literature ^26^ for Smc1, a subunit of the cohesin complex, to analyse cohesin localisation before and after Spt16 depletion along yeast chromosomes (Fig. 2a). Centromeric regions are ideal sites to study cohesin translocation because cohesin complexes are initially loaded at core *CEN* sequences and then move away towards pericentromeric regions ^15^. We observed that cohesin localisation to core *CEN* sequence did not change upon Spt16 degradation, however its accumulation at pericentric regions was severely reduced (Fig. 2a-b). This suggest that FACT might not be required for cohesin initial loading at *CEN* sites ^26^, but it is necessary for its accumulation on peri-centric regions (Fig. 2b). Mcd1 is expressed at low levels in G_1_-arrested cells, thus cohesin association to pericentric regions is very modest at this stage of the cell cycle ^15^. However, cohesin loading can still occur in G_1_ ^26^ when cohesin expression is artificially induced. Indeed, expression of *MCD1* from the *GAL1-10* promoter caused a significant enrichment of Smc1 both at core *CEN* sequences and the surrounding pericentric regions (Fig. 2c). In contrast, *MCD1* expression in G_1_-arrests depleted in Spt16 showed no enrichment of cohesin at pericentric regions (Fig. 2c), despite the fact that Smc1 still was present at core *CEN* sequences (Fig. 2c). Therefore, these results demonstrate that in the absence of the histone chaperone FACT, cohesins can be loaded at core *CEN* sites but fail to accumulate at pericentric regions.

**Figure 2.**
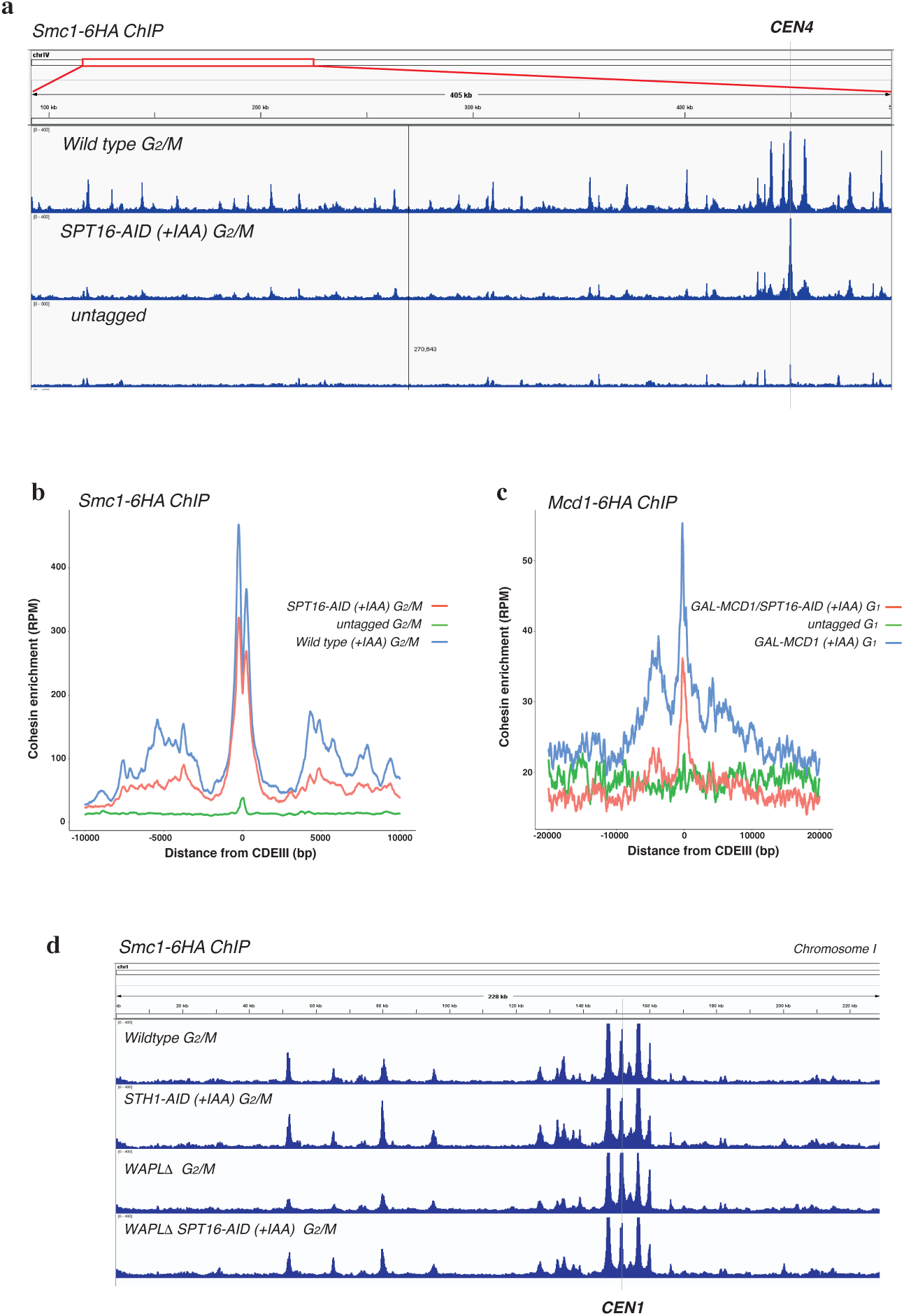
FACT is necessary for the localization of cohesins in metaphase chromosomes. **a**, Calibrated ChIP-seq in strains carrying either *SMC1-6HA or SMC1-6HA SPT16-AID* (which allows auxin-mediated degradation of Spt16). Cells were arrested in metaphase (G_2_/M-nocodazole) and exposed to auxin to degrade Spt16. Western analysis for Spt16 confirmed the degradation of Spt16 in *SPT16-AID* cells upon auxin addition (Suppl. Fig. 1). Nuclear and cellular morphology was used following auxin addition to confirm that the cell population remained arrested in metaphase (Suppl. Fig. 1). Smc1-6HA calibrated ChIP-seq signal profile along a 150Kb region of yeast chromosome 4 in *SMC1-6HA, SMC1-6HA SPT16-AID* and untagged strains are shown. **b**, Calibrated ChIP-seq comparing average profile of *SMC1-6HA* around *CENs* in G_2_/M (nocodazole) in *SMC1-6HA, SMC1-6HA SPT16-AID* or untagged strains as in **a**. The number of reads at each base pair from CDEIII was averaged over all 16 chromosomes. **c**, Calibrated ChIP-seq comparing average profile of *MCD1-6HA* around *CENs* in cells arrested in G_1_ (*α*-factor) overexpressing *MCD1* from the galactose promoter, in the presence or absence of Spt16 (*SPT16-AID*). The number of reads at each base pair from CDEIII was averaged over all 16 chromosomes. **d**, Calibrated ChIP-seq in strains carrying either *SMC1-6HA or SMC1-6HA STH1-AID* (which allows auxin-mediated degradation of Sth1), *SMC1-6HA WAPIΔ* and *SMC1-6HA WAPIΔ SPT16-AID.* Cells were arrested in metaphase (G_2_/M-nocodazole) and exposed to auxin to degrade Sth1 or Spt16. Smc1-6HA calibrated ChIP-seq signal profile along yeast chromosome 1 in *SMC1-6HA, SMC1-6HA STH1-AID, SMC1-6HA WAPIΔ*, *SMC1-6HA WAPIΔ SPT16-AID* and untagged strains are shown.

To confirm that the loss of cohesin signal in FACT-depleted cells was not due to a defect in cohesin loading, we first tested whether cohesin loading is high in G_2_/M arrested cells (the stage where we depleted FACT in our experiments). To do this, we used an auxin (IAA)-inducible degron of Sth1 (*STH1-AID*), a core subunit of RSC, because RSC is required for cohesin loading^6^. Cohesin binding around centromeres and chromosome arms was similar to wildtype cells in the Sth1-depletion (Fig. 2d & Suppl. Fig. 2), demonstrating that *de novo* cohesin loading in metaphase arrested cells is likely to be very low. As a second approach to rule out a role for FACT in cohesin loading, we tested whether cohesin loaded when FACT is depleted in the absence of the cohesin release factor Wapl ^27^. We observed that cohesin levels on chromosomes were not reduced in *WAPLΔ SPT16-AID* cells (Fig. 2d), confirming that the loss of cohesin signal in FACT-depleted cells cannot be caused by an impairment in cohesin loading but rather the loss of stability of cohesin on chromatin.

In yeast, cohesin complexes can be divided into two classes with respect to their chromosomal association mechanisms, those loaded at *CEN* sequences, which move and accumulate in pericentric regions ^28,29^, and those that load at chromosomal arm sites and translocate to regions of convergent transcription ^30^. Importantly, the translocation of cohesins from *CEN* sites to pericentromeric regions requires Scc2 ^15^ and is independent of transcription ^15^, while the movement of arm loaded cohesins to sites between convergent genes involves relocation dependent on transcription ^30-33^.

FACT plays important roles during transcription initiation and elongation^34,35^. Therefore, we decided to investigate to what extend changes in transcription upon FACT-depletion are responsible for the disruption in the cohesin ChIP profiles. First, we focused on centromeric regions where transcription should not affect movement of *CEN* loaded cohesins to pericentric regions ^15^. To confirm that accumulation of cohesin at pericentromeric regions does not require active transcription, we treated wildtype cells with the transcription repressor thiolutin and used calibrated ChiP-seq to Smc1 to compare cohesin enrichment in the presence and absence of active transcription (Fig. 3a). Thiolutin treatment had little effect on Smc1 binding at core *CEN* sequences or pericentric sites (Fig. 3a). Therefore, as reported previously^15^, accumulation of cohesin at these sites is independent of transcription. In contrast, Spt16 degradation in thiolutin-treated cells caused a reduction in cohesin presence on pericentric regions (Fig. 3a). Therefore, at pericentric regions FACT facilitates cohesin accumulation independently of active transcription.

**Figure 3.**
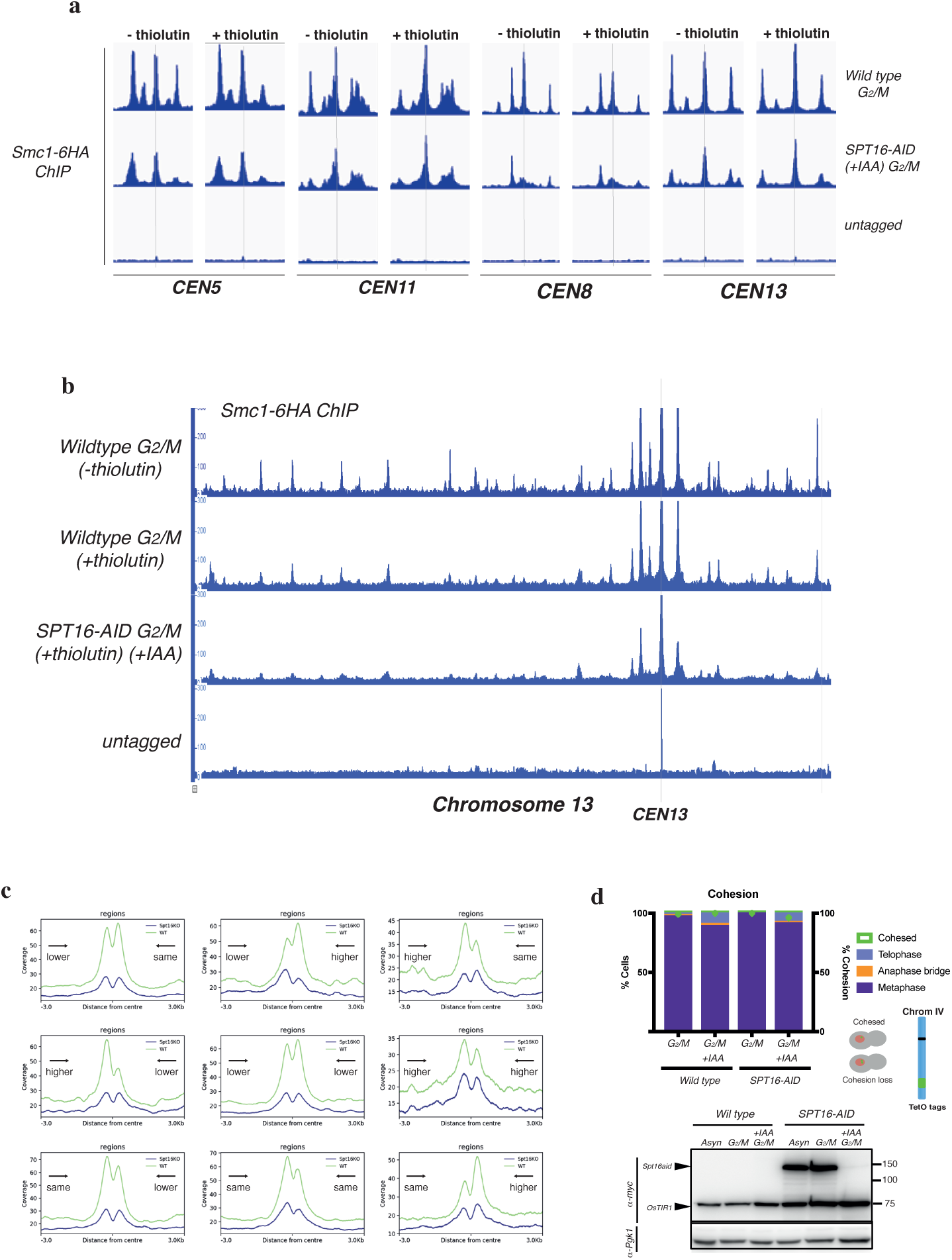
FACT inactivation causes defects in chromosome organisation but not sister chromatid cohesion. **a**, Calibrated ChIP-seq profiles of *SMC1-6HA* around *CEN5, CEN8, CEN11* and *CEN13* in G_2_/M (nocodazole) in *SMC1-6HA, SMC1-6HA SPT16* and untagged strains in the presence and absence of the transcription inhibitor thiolutin. Cells were arrested in G_2_/M metaphase and exposed to auxin to degrade Spt16 (in *SPT16-AID*). Half of the culture was treated with thiolutin (+thiolutin) to block transcription. **b**, Calibrated ChIP-seq profiles of *SMC1-6HA* along yeast chromosome 13 in G_2_/M (nocodazole) in *SMC1-6HA, SMC1-6HA SPT16* and untagged strains in the presence and absence of the transcription inhibitor thiolutin as indicated (+/- thiolutin). **c**, Density profiles of Smc1 binding at convergent gene sites in *SMC1-6HA, SMC1-6HA SPT16* strains. Cells were arrested in G_2_/M (nocodazole) and exposed to auxin to degrade Spt16 (in *SPT16-AID*). Convergent gene sites were subdivided according to transcriptional activity changes upon Spt16 depletion. **d**, Analysis of sister chromatid cohesion in the absence of FACT. *SPT16-AID* and control cells were arrested in metaphase and exposed to auxin to degrade Spt16. Protein levels (westerns) and cell cycle stage analysis for the experiment are shown (right graph). The strains carried tetO/tetR-based chromosome tags inserted at the right subtelomere region of chromosome IV that were used to score loss of cohesion (right graph). Note that auxin addition leads to the degradation of Spt16 in *SPT16-AID* cells (westerns) and the metaphase arrest is stable throughout the experiment. Cohesion loss was not increased in the absence of Spt16.

The analysis of cohesin translocation on chromosome arms is more complicated than at centromeres because it is difficult to detect cohesin during loading, and consequently ChIP profiles at arm regions reflect the final position of the complexes. Since our results demonstrate that FACT inactivation also causes a decrease in cohesin binding at chromosomal arm sites (Fig. 2a). Nevertheless, we investigated whether FACT plays a role in cohesin stability independent on transcription. Smc1 binding across chromosome arm sites was reduced in wildtype cells treated with thiolutin (Fig. 3b) confirming that active transcription is important for cohesin enrichment at these sites ^30-33^. Thiolutin treatment in FACT-depleted cells caused a further reduction in cohesin binding across chromosome sites (Fig. 3b).

To further dissect the contribution of FACT to cohesin stability, we analysed how Smc1 enrichment at convergent transcription sites correlates with the changes in transcription at these sites when Spt16 is depleted (Fig. 3c). Analysis of cohesin binding sites confirmed their enrichment in positions in the genome where transcription units are oriented in a convergent manner ^30-32^. Interestingly, cohesin enrichment was observed as a double peak at both 3’ ends of the genes (Fig. 3c). We classified convergent sites into categories depending on whether the transcription of the flanking genes had change by 1.5-fold up or down, or had remain the same (Fig. 3c) upon Spt16 depletion. We then compared cohesin enrichment changes in each transcription-pair category (Fig. 3c). Smc1 was reduced similarly in all categories (Fig. 3c). Therefore, we conclude that FACT has a direct function in cohesin stability on chromosome arm sites and that changes in cohesin binding during cannot be explained solely by the altered transcription profiles upon depletion of Spt16.

Having demonstrated that FACT has a role in maintaining cohesin on chromosomes (Fig. 3a-c), we sought to investigate the functional consequences for chromosome architecture. Cohesin holds sister chromatid together ^1,2^ and organises intra-chromatid loops that provide an integral level of structure to chromosomes ^4,5^. First, we tested whether cohesion between sister chromatids is affected in G_2_/M arrested cells when Spt16 is depleted. We scored cohesion using tetO/tetR-based tags at sub-telomeric regions of chromosome IV (Fig. 3d). Surprisingly, cells depleted for Spt16 showed no defects in cohesion (Fig. 3d). Next, we tested whether Spt16 is necessary to maintain mitotic chromosome architecture. Recent studies on mitotic yeast chromosomes using Hi-C have shown that cohesin compacts chromosome arms through *cis* interactions, independently of sister chromatid cohesion ^4,5^. We therefore considered the possibility that the reduction in cohesins observed upon Spt16-depletion (Fig. 2a) might impact on *cis* contacts along yeast metaphase chromosomes. To explore this, we built Hi-C libraries from G_2_/M arrested cells depleted for Spt16 and compared it to control arrests (Fig. 4a and Suppl. Fig. 3). The resulting contact maps show that depletion of Spt16 led to a reduction in short-distance intra-chromosomal contacts while long-range intra-chromosomal contacts were increased (> 100kb), as suggested by i) a larger width of the main diagonal in the contact maps of cells depleted for Spt16 (Suppl. Fig. 3), and as confirmed by computing ii) the contact probability p as a function of genomic distance s of all chromosome arms (Fig. 4b) and iii) the log-ratio between the normalized contact maps of depleted and non-depleted cells (Fig. 4a). The ratio map also shows that Spt16 depletion cause an increase of inter-chromosomal contacts (Fig. 4a). These results suggest that chromatin fibers loose internal organization in the absence of Spt16.

**Figure 4.**
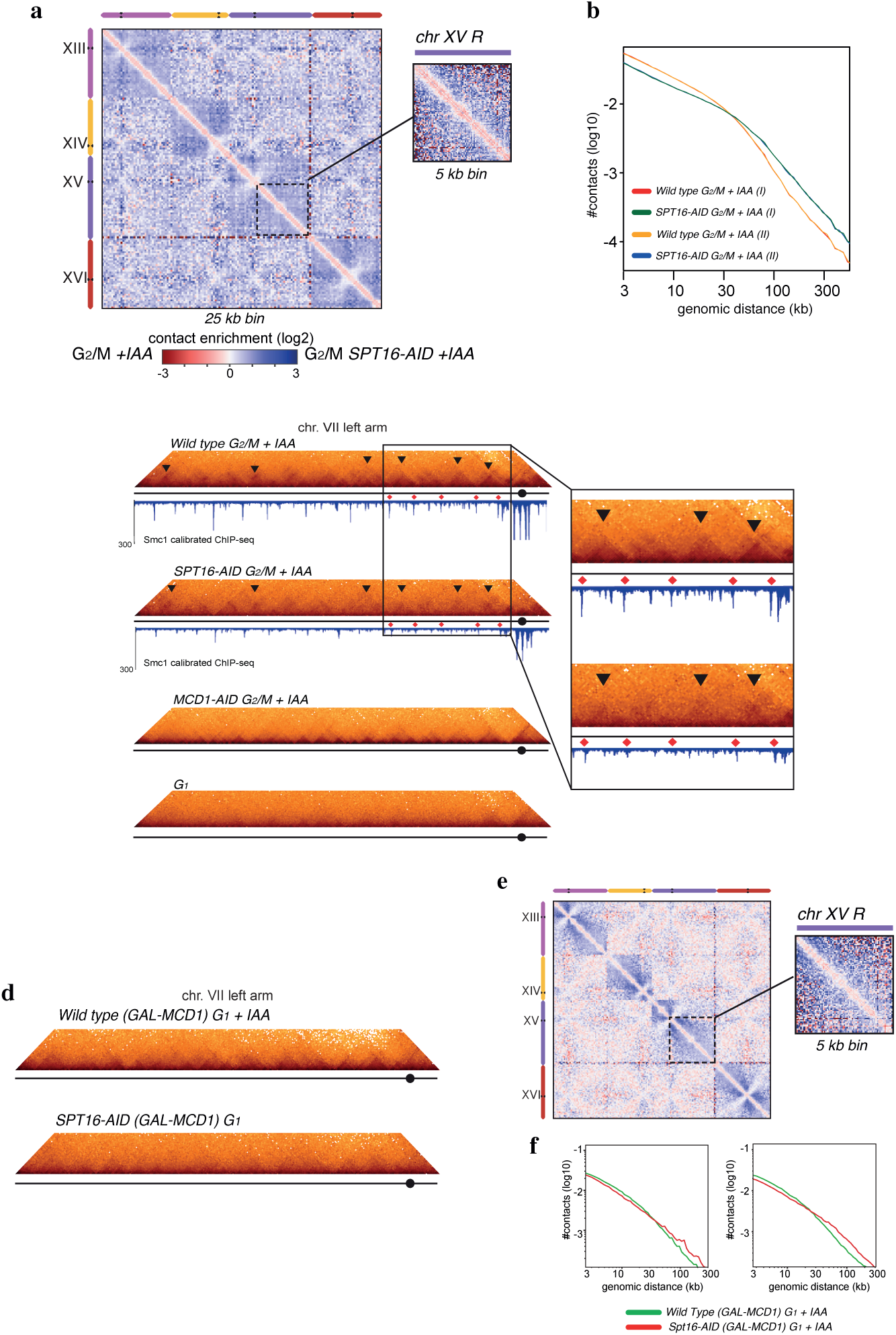
FACT function is important for the establishment cohesin-dependent TAD-like domains. **a**, Log_2_ ratio of the normalized contact map (25 kb bins) of Spt16-aid (spt16 depleted) vs. wild-type (top right) metaphase arrested cells. Blue to red color scales reflect the enrichment in contacts of wild-type versus spt16-aid respectively. Inset display magnifications of chr7 at 25 and 5 kb resolution. **b**, Contact probability (P(s)) for wild-type and spt16-aid (spt16 depleted) metaphase cells averaged over all chromosomes is shown. **c**, Magnification of the left arm of chromosome VII normalized contact maps (2 kb bins) of G_1_ (Wild type G_1_) and G_2_/M arrested cells in the presence (Wild type G_2_/M+IAA) and absence of Spt16 (*SPT16-AID* G_2_/M+IAA) or Mcd1 (*MCD1-AID* G_2_/M+IAA). The plot under map represents the enrichment in cohesin at this stage as quantified by calibrated ChIP-seq. Black triangles point at TAD-like signals that are lost upon Stp16 depletion. Red diamonds: enrichment peaks between the TAD-like signals that are lost upon Spt16 depletion. Inset display magnifications of chr7 pericentromeric regions where cohesin sites (red diamons) and TAD-like domains are lost in Spt16 depleted cells compared to control G_2_/M arrested cells. **d**, Magnification of the left arm of chromosome VII normalized contact maps (2 kb bins) of G_1_ arrested cells overexpressing *MCD1* from the galactose promoter in the presence (Wild type *GAL-MCD1* G_1_+IAA) or absence (*SPT16-AID GAL-MCD1* G_1_+IAA) of Spt16. **e**, Log_2_ ratio of the normalized contact map (25 kb bins) of G1 arrested cells overexpressing MCD1 in the presence and absence of Spt16. Blue to red color scales reflect the enrichment in contacts of wild-type versus Spt16-depleted arrests respectively. Inset display magnifications of chr7 at 25 and 5 kb resolution. **f**, Contact probability (P(s)) for G_1_ arrested cells overexpressing *MCD1* in the presence (Wild type *GAL-MCD1* G_1_+IAA) and absence (*SPT16-AID GAL-MCD1* G_1_+IAA) of Spt16. Duplicate experiments are shown.

To further characterize mitotic chromosomes, 2 kb contact maps were generated. At this resolution, triangular contact patterns reminiscent of the topological associating domain (TADs) described in metazoans appear along mitotic chromosomes in G_2_/M arrested cells but not G_1_ arrests (Fig. 4c). The boundaries of these domains involve regions enriched in cohesin (red diamonds, Fig. 4c), and stronger contacts between pairs of boundaries are often visible, consistent with the establishment of cohesin loops (black triangles, Fig. 4c) (previous Hi-C studies on cohesin failed to directly characterize these loops, as the experiments were not analyzed at this resolution ^4,5^). We tested whether the triangular patterns of the 2 kb contact maps were dependent on cohesin. To this aim, we built Hi-C libraries from cells depleted for Mcd1 while arrested in G_2_/M (Fig. 2c). The resulting contact maps showed a significant reduction in TAD-like structures (Fig. 4c), as expected from previous reports demonstrating a key role for cohesin in loop formation ^3-5^. The depletion of Spt16 results in a strong reduction of cohesin deposition (Fig. 4c), as well as a fading of the cohesin-dependent TADs-like domains signal (Fig. 4c).

Our Hi-C analysis is consistent with a role for FACT in affecting cohesin-dependent loops, however we cannot differentiate whether FACT is important to establish these loops or to maintain them. To test a potential role for FACT in *de novo* loop formation by cohesin, we first investigated whether overexpressing *MCD1* in G_1_, which causes cohesin loading (Fig. 2c), leads to *de novo* formation of loop, which are normally absent in G_1_ cells ^4,5^ (Fig. 4c). Hi-C libraries from G_1_ arrested cells overexpressing Mcd1 from the *GAL1-10* promoter were thus generated. TADs-like domain signals were observed in 2 kb contact maps of G_1_ overexpressing Mcd1 (Fig. 4d) demonstrating *de novo* loop formation by cohesin under this experimental condition (Fig. 4d). Next, we depleted Spt16 during the overexpression of Mcd1 in G_1_ cells, cohesin dependent loops were now strongly reduced, and hardly visible, in the absence of FACT (Suppl. Fig. 4). The log-ratio between the normalized contact maps of G_1_ depleted and non-depleted cells (Fig. 4e) and the contact probabilities as a function of genomic distance s of all chromosome arms (Fig. 4f) confirms that *de novo* loop formation by cohesins in G_1_ requires FACT.

Our results demonstrate that while cohesin’s role in sister chromatid cohesion is maintained in the absence of FACT (Fig. 3d), its function in organising chromatin loops in metaphase chromosomes is impaired (Fig. 4a-c). We therefore hypothesized that depletion of FACT in metaphase arrested cells might lead to defects in chromosome structure that are likely to compromise the ability of chromosomes to segregate faithfully if allowed to proceed into anaphase. To test this, we arrested *SPT16-AID* cells in metaphase, depleted FACT, and released cells from the arrest. To ensure that cells did not pass beyond telophase, so that we could accurately score chromosome segregation, we performed these experiments in a *CDC15-AID* genetic background. Therefore, upon release from metaphase, the addition of auxin to degrade Spt16, also induced the degradation of the mitotic exit kinase Cdc15, causing a tight telophase arrest. Similar to our analysis of sister chromatid cohesion, we used tetO/tetR-based tags on chromosome IV to evaluate chromosome segregation in the Cdc15 block. As predicted, we observed high levels of chromosome missegregation in cells depleted for Spt16 and Cdc15, but not Cdc15 only (Suppl. Fig. 5).

In summary, our work demonstrates that the FACT complex is a mediator of cohesin stability on chromatin (Fig. 2b-c) and through this function facilitates the formation of cohesin-dependent loops (Fig. 4). Recently, an involvement of FACT in chromatin architecture independent of transcription has been proposed ^36^, our data suggests that this function might be related to cohesin’s role in genome architecture. We speculate that similar to its roles in transcription and replication, FACT function as a histone chaperone is likely to occur through destabilization of core nucleosomal particles, to allow movement of cohesins on chromatin. We demonstrate that FACT contributes to cohesin’s role in organizing mitotic chromosome folding but not sister chromatid cohesion. Finally, our results support the original claim that cohesin complexes have two distinct roles, in chromosome organisation and sister chromatid cohesion ^1 37^ and demonstrate that FACT is necessary for cohesin functions in genome folding.

## Methods

Full detail of methods used in this study can be found in the Supplementary materials section. For gel source data, see Supplementary Figures.

## Acknowledgements

We thank Axel Cournac for analyses; and the members of the L.A, R.K. and P.A. laboratories for fruitful discussions and advice. The work in the L.A. laboratory was supported by Wellcome Trust Senior Investigator award to L.A. (100955, “Functional dissection of mitotic chromatin”) and the “London Institute of Medical Research (LMS), which receives its core funding from the UK Medical Research Council. This research was further supported by funding from The European Research Council (R.K.), Agence Nationale pour la Recherche (R.K.) and the the Spanish Ministerio de Economía, Industria y Competitividad (BFU2017-89622-P) (F.A.).

