## Supplementary materials and Figures for "FACT mediates cohesin function on chromatin"

#### Methods

##### Yeast strain and DNA constructs

Yeast strains used in this study are listed in Table 1. Genetic background of these strains is YPH499 (congenic of S288C) except for (mass spec strains) that is BY4741. Epitope tagging of genes and deletions were carried out as described [1].

| | | |
|---|---|---|
| *CCG4000* | *MATa ura3-52 lys2-801 ade2-101 trp1-Δ63 his3-Δ200 leu2-Δ1 bar1-Δ* | *A. Strunnikov* [a] |
| *CCG12978* | *CCG4000 ADH1-OsTIR1:9Myc(URA3) PMETCDC20 (TRP) Scc1:AID*-9myc:HphNT1* | *This study* |
| *CCG14758* | *CCG4000 ADH1-OsTIR1:9Myc(URA3) SMC1:6HA:NatNT2* | *This study* |
| *CCG14759* | *CCG14758 Spt16:AID*-9myc:KanMX* | *This study* |
| *CCG14762* | *CCG4000 SMC1:6ha:NatNT2* | *This study* |
| *CCG14763* | *CCG14762 SPT16:9myc:HphNT1* | *This study* |
| *CCG13889* | *CCG4000 ADH1-OsTIR1:9Myc(URA3) TetR-GFP (ADE2) PMETCDC20 (TRP) TetO:1513Kb ChrIV (HIS3) cdc15:AID-9myc:HphNT1* | *This study* |
| *CCG14764* | *CCG13889 Spt16:AID*-9myc:KanMX* | *This study* |
| *CCG14767* | *MATa his3Δ1 leu2Δ0 met15Δ0 ura3Δ0 NAT::GAL-CDC20* | *This study* |
| *CCG14768* | *MATa his3Δ1 leu2Δ0 met15Δ0 ura3Δ0 SCC1-6HA::HPH NAT::GAL-CDC20* | *This study* |
| *CCG13142* | *Mata leu2 ura3 his3 trp1 ade2 lys2 bar1 pep4:HIS3 ; pADH1-OsTIR1::LEU* | *This study* |
| *CCG14795* | *CCG13142; Gal-Scc1-6HA::ADE* | *This study* |
| *CCG14796* | *Mata leu2 ura3 his3 trp1 ade2 lys2 bar1 pep4:HIS3 ; pADH1-OsTIR1::LEU; Spt16-AID-9myc::Kan; Gal-Scc1-6HA::ADE* | *This study* |
| *CCG14696* | *Mata leu2 ura3 his3 trp1 ade2 lys2 bar1 pep4:HIS3 ; pADH1-OsTIR1::LEU; Sth1-AID-9myc::Nat; Smc1-6HA::KanMX* | *This study* |
| *CCG14780* | *Candida glabrata Scc1-3HA* | *K. A. Nasmyth* [15] |

[a] Freeman, L., AragonAlcaide, L. & Strunnikov, A. The condensin complex governs chromosome condensation and mitotic transmission of rDNA. *J. Cell Biol.* **149**, 811–24 (2000).

**Media, culture conditions and synchronisations**

To arrest the cells in G1, α-factor was added to exponentially growing MATa cultures (OD600=0.5) to a final concentration of 3 x 10-8 M. To arrest cells in G2/M, Nocodazole (1.5 mg/ml stock in DMSO 100%) was added to exponentially growing cultures (OD600=0.5) to a final concentration of 0.015 mg/ml. When arrests in Nocodazole are maintained for more than 2 hours cells are spun and resuspended in YPD containing fresh nocodazole (0.015 mg/ml). Cultures were monitored by microscopy until ≥90% of cells were arrested, typically 2 hours at 25 °C. To release cells from G1, the culture was spun (4,000 r.p.m, 1 min) and washed in YPD 3 times. The pellet was then resuspended in YPD containing 0.1 mg/ml pronase. To release cells from Nocodazole, the culture was spun (4,000 r.p.m, 1 min) and washed in YPD containing 1% DMSO 5 times. The pellet was then resuspended in YPD. To degrade proteins tagged with AID* epitope, a stock of IAA of 0.6M in ethanol 100% was used for a final concentration of 6mM. Where removal of IAA from the media was carried out, cells where spun (4,000 r.p.m, 1 min) and washed 5 times in YPD. The pellet was then resuspended in YPD. To inhibit transcription a stock of thiolutin at 1mg/ml in DMSO was added for 30 minutes to the culture at a final concentration of 10ug/ml.

**Microscopy**

A Leica IRB epifluorescence microscope with a Hamamatsu 742-95 digital camera, a 63X/1.4 lens and OpenLab software (Improvision) was used for single cell visualization. 1 ml of cell culture was collected from each time point and mixed with glycerol (20% final concentration) to preserve TetO/TetR signal. Cells were then frozen at -80 °C. For visualization, cells where centrifuged at 3,000 r.p.m. for 2 minute and 1μl of the pellet was mixed with 1μl of DAPI solution (DAPI 4 μg/ml Triton 1 %) on the microscope slide. For each field 20 z-focal planes images were captured (0.3 μm depth between each consecutive image). At least 100 cells were score for GFP dots.

**Hi-C libraries**

Hi-C experiments were performed according to a protocol adapted from [2]. Aliquots of 1-3 x $10^9$ cells in 150 ml culture medium were fixed in 3% formaldehyde (Sigma-Aldrich, Ref. F8775) and quenched in 0.4 M glycine for 20 min at room temperature and 4°C, respectively. Cells were harvested, washed with culture medium and pellets were stored at -80°C. Pellets were thawed on ice and resuspended in chilled TE buffer supplemented with protease inhibitors. Cells were lysed using Precellys VK05 Lysing KIT (Ozyme, Ref. 0042) at 6,700 rpm. Lysates were treated with 0.5 % SDS for 20 min at 65°C and digested overnight with DpnII (CFinal=500 U/pellet; NEB R0543M) at 37°C. Digestions were centrifuged for 20 min at 16,000 g and the pellets suspended in cold water. 5' DpnII overhangs were filled-in and labelled with biotin using 30 µM dNTP (dATP, dGTP, dTTP and Biotin-14-dCTP Invitrogen Ref. 19518018). Biotinylated DNA fragments were ligated using T4 DNA ligase (CFinal=250 Weiss U/pellet; Thermo Scientific 10621441) for 4 h at 16°C. Cross-link was reverted through an overnight treatment in presence of 250 µg/ml proteinase K at 65°C. Total DNA was extracted using phenol/chloroform, precipitated and treated with RNAse A 500 µg/ml. Non-ligated DNA fragments were removed by T4 DNA polymerase (CFinal=5 U/pellet; NEB M0203L) treatment; whereas ligated fragments were 500 bp sheared using Covaris S220 and pulled-down using Dynabeads Myone Streptavidin C1 (Invitrogen 65001). Resulting libraries were amplified using custom-made primers for paired-end deep-sequencing on NextSeq500 Illumina platform (2x75 bp cycles).

**Generation and normalization of contact maps**

The reads from each Hi-C library were processed as follows. First, PCR duplicates were removed using the custom-made tags present on the primers and the resulting reads were mapped independently using Bowtie 2 (mode: --very-sensitive --rdg 500,3 --rfg 500,3) against the reference genome of S. cerevisiae (S288C) and indexed on DpnII restriction fragments. An iterative alignment procedure (20 bp window) was used to maximize the yield of uniquely mapped reads (mapping quality > 30). Reads were classified as either valid Hi-C products or unwanted events to be filtered out (i.e. loops, non-digested fragments, etc.; for details see {Cournac, 2012 #40} ) based on their assignment and orientation on the DpnII restriction fragments in the reference genome. Valid Hi-C reads were used to generate contact frequency maps. In these maps, each vector consists in a fixed size genomic bin made of successive RF (either 5, 25, or 50 kb). Bins with a high variance in contact frequency (< 1.5 Standard Deviation or 1.5 - 2 S.D.) were filtered out. Filtered contact maps were normalized as described [3]. Approximately 15 million of valid reads were used to build each contact map. Similarity between pairs of normalized contact maps was assessed by computing the log2 ratio for each point.

**Contact probability as a function of the genomic distance p(s)**

The contact probability p(s) as a function of the genomic distance s separating two DNA positions in cis was computed as follow. First, self-circularization events were discarded by removing pairs of read oriented in opposite directions along the reference genome and separated by less than 1.5 kb. The remaining reads were log-binned as a function of their genomic distance (in kb) on each chromosomal arm: bin = [log1.1(s)]. The p(s) curve is the histogram computed on the sum of read pairs for each bin. This sum is weighted by both bin-size $1.1^{(1+bin)}$ (log binning), as well as by the length of the chromosomes.

**Chromatin immunoprecipitation**

For ChIP analysis cells were grown to $OD_{600}$ = 0.5 and arrested at the required cell cycle stage. 87 $OD_{600}$ units of *S. cerevisiae* were mixed with 43 $OD_{600}$ units of asynchronous *C. glabrata*. Cells were fixed for 15 minutes at 25 °C, and quenched with glycine (final concentration 125 mM) for 7 minutes before cells were harvested by centrifugation at 4,000 r.p.m. for 1 minute. The cell pellets were washed in PBS and transferred to a screw cap tube and frozen on dry ice. The pellets were stored at -80 °C. Pellets were resuspended in 300 µl of IP buffer (150 mM NaCl, 50 mM Tris-HCl (pH 7.5), 5 mM EDTA, NP-40 (0.05% v/v), Triton® X-100 (1% v/v)) containing phenylmethanesulfonyl fluoride (PMSF, final concentration 1 mM) and Complete protease inhibitor cocktail (without EDTA, from Roche). 500 µl of glass beads were added to the tubes. Cells were broken in a FastPrep® FP120 (BIO101) by 3 repetitions of a 20 seconds cycle, power setting 5.5. Cells were maintained on ice for 2 minutes after each cycle. The tubes were pierced with a hot needle and placed into new eppendorf and spun at 2000 r.p.m. for 2 minutes to collect the lysate. 100 µl of IP buffer containing PMSF and protease inhibitors were then added and tubes spun again at 2000 r.p.m. for 1 minute. The cell lysate was spun down for 10 minutes at 15,000 r.p.m at 4 °C. This pellet was resuspended in 1ml of IP buffer containing PMSF and protease inhibitors, and sonicated for 1 hour (30 seconds on, 30 seconds off) at high power at 4 °C in a Diagenode Bioruptor. After sonication samples were spun down for 10 minutes at 15,000 r.p.m. and the supernatant taken. 200 µl of the sonicated chromatin were taken as “input” and 400 µl was incubated with 40 µg of HA antibody (anti-HA 12CA5 from Roche) in a sonicator at low power for 30 minutes (30 seconds on, 30 seconds off). The “input” DNA was precipitated with 0.3 M sodium acetate and 2.5 volumes of cold ethanol and spun down at 15,000 r.p.m. for 30 minutes, and the supernatant removed. The pellet was washed with 70 % ethanol and air dried. After antibody binding the IP sample was spun down at 13,000 r.p.m for 5 minutes and the supernatant added to 60 µl of Dynabeads protein G (Invitrogen) previously equilibrated with IP buffer. Then the samples were incubated for 2 hours at 4 °C in a rotating wheel and washed 5 times with IP buffer using a magnetic separator rack. Finally “input” samples and IP samples were resuspended in de-crosslinking buffer (TE 1X, 1 % SDS, 10 μg/ml RNase A , 1 mg/ml proteinase K) and incubated at 65 °C overnight. Samples were purified using ChIP DNA Clean & Concentrator kit (Zymoresearch) according to the manufacturer instructions.

**Protein coimmunoprecipitation**

Cells were grown to $OD_{600}$ = 1 wash in cold water and resuspended in ice-cold buffer (50 mM HEPES, 150 mM KCl, 1.5 mM MgCl2, 0.5 mM DTT, and 0.5% Triton X-100 (pH 7.5) supplemented with Complete protease inhibitor cocktail tablets (from Roche)). Cells were broken in a FastPrep® FP120 (BIO101) by 3 repetitions of a 20 seconds cycle, power setting 5.5. Extracts were maintained on ice for 2 minutes after each cycle. To the cell extract 0.5mM $CaCl_2$, 20 U DnaseI (NEB), 0.2 mg RNaseA (Qiagen) and 500 U Benzonase (Millipore) were added in order to degrade DNA and RNA from the sample. Cell extracts were incubated with protein G Dynabeads (Invitrogen) bound to anti-Myc antibody (Roche, 9E10) for 2 h at 4°C. Finally, beads were washed 5 time in washing buffer (10mM Tris-Cl pH 7.5, 150mM NaCl, 0.5 % Triton) and unbound from the antibody by incubating at °C for 4 minutes in SR Buffer (2% SDS, 0.125 M Tris-Cl pH 6.8). Immunoprecipitated proteins were mixed with SS buffer (5% sacarose, 0.0125 % Bromophenol blue) and run in SDS-PAGE gel.

**Digestion with MNase**

Mononucleosomal DNA was prepared as described in [4]. For MNase digestions 160 ml of cells (OD600=1) where fixed with 1 % formaldehyde for 30 minutes at 25 °C with 100 r.p.m. agitation and quenched with 125 mM glycine for 10 minutes at 25 °C with 100 r.p.m. agitation. Cells were spun and washed twice with PBS. Pellet was resuspended in 20 ml sorbitol/ tris buffer (1M sorbitol, 50 mM Tris-HCl pH 7.5, 10 mM β-mercaptoethanol) containing 10 mg of Zymolyase 20T and incubated at 30 °C for 5 minutes or until spheroplast were formed. Spheroplasts were spun at 3,000 r.p.m. for 5 minutes and washed with sorbitol/tris buffer without β-mercaptoethanol. Spheroplasts were resuspended in NP buffer (1 M sorbitol, 50 mM NaCl, 10mM Tris pH 7.5, 5 mM MgCl2, 1mM CaCl2, 1mM □-mercaptoethanol, 500 μM spermidine, 0.075 % NP40) and digested with 600, 1200 or 1800

U/ ml of MNase (NEB) for 20 minutes at 37 °C. MNase digestion was stopped adding 200 µl of STOP solution ( 5% SDS, 50 mM EDTA). DNA was decrosslinked by adding 1 mg/ml proteinase K and incubating at 65 °C overnight. DNA was purified by adding 0.6M potassium acetate pH 5.5 and incubating on ice 5 minutes and centrifuging 10 minutes at 3,500 r.p.m followed by a phenol chloroform extraction. Finally DNA was precipitated by adding 25 µg/ml glycogen, 1/25 columes NaCl, 0.7 volumes Isopropanol, incubating 30 minutes at -20 °C and centrifuging 30 minutes at 13,200 r.p.m. The pellet was washed in 70 % ethanol and resuspended in TE 1X. Samples were run in parallel in a 1.5 % agarose gel and the mononucleosomes of a digestion with a ratio of 80:20 mononucleosomes to dinucleosomes were purified and used in Next Generation Sequencing.

**Next-generation sequencing and DANPOS analysis**

Libraries of mononucleosomal and ChIP DNA were constructed following the Illumina compatible NEXTflexTM ChIP-Seq kit protocol and sequenced in an Illumina NextSeq500 platform using the paired-end protocol. Reads were aligned to the S.cerevisiae (SacCer3) genome using Bowtie [5]. Nucleosome occupancy maps were generated using the NUCwave algorithm [6]. Nucleosome fuzziness was analyzed using the dpos utility of DANPOS2 application [7]. To make it compatible with NUCwave maps, we used the following parameters: a span of 1 bp and a read extension of 50 bp. ChIP-seq data representation was performed using the Model-based Analysis of ChIP-Seq (MACS) [8] with a read extension of 147 bp.

**Calibrated ChIP-Seq analysis**

Calibrated Chip-seq analysis as done as in [9]. The quality of the raw sequence data was assessed using FastQC (version 0.11.5). The reads were mapped to the S. cerevisiae genome (sacCer3) as the experimental genome or C. glabrata genome (CBS138) as calibration genome using Bowtie2 (version 2.3.4) with the default parameters. To generate an alignment in which all the sequences exclusively map to the sacCer3 genome, the whole reads were first mapped to CBS138 genome and the unmapped reads were retrieved as a separate Fastq file. Subsequently, these unmapped reads were re-aligned to sacCer3 genome and the generated aligned BAM file therefore contained reads that were unique to sacCer3. A same method was used to generate alignments unique to CBS138.

Applying a recent published method [26] for calibrating ChIP-seq data and adding cells with the calibration genome (CBS138) to different samples of cells with the experimental genome (sacCer3), we calculated occupancy ratio (ORi) for experimental sample i from the number of reads assigned to calibration (WCi) and experimental (WXi ) genomes among sequences derived from input DNA and the number of reads assigned to calibration (IPCi ) and experimental (IPXi) genomes among sequences derived from ChIP-seq sample. Then ORi was calculated as:

$$ORi = \frac{Wci}{Wxi} \times \frac{IPxi}{IPci}$$

For each sample, the alignment BAM file was converted to Wig format using bedtools (version 2.27.1). To obtain a calibrated version, the Wig file with the conventional ChIP-seq profile was multiplied by the occupancy ratio (OR) corresponding to each sample. Subsequently, these calibrated BigWig files were used to generate average profile plots for regions in +/- 20KB of centromeres for 16 chromosomes in sacCer3 genome
All sequencing data that support the findings of this study have been deposited in GEO and are accessible through the following accession number: GSE118534.

**MNase-seq data analysis**

Paired-end Sequencing reads were aligned to Yeast genome sacCer3 using bowtie version 2.2.9 with default parameters. DANPOS2 was applied for two separate pair-wise comparisons at each nucleosome between G1 arrest and G2/M arrest, and G2/M arrest and G2/M +IAA. DANPOS output for pair-wise comparison contains a set of reference positions (with fixed length of 200bp) for nucleosomes defined by integrating all the nucleosomal positions called in each of the two conditions under study. The DANPOS-generated reference positions in two pair-wise comparisons were very similar but there was a small fraction of positions that did not overlap well in two comparisons. DANPOS also calculated Fuzziness scores for each condition and FDR and log10 p-value for each pair-wise comparison. We reported the nucleosomes with significant fuzziness change as reference nucleosomes in each comparison with FDR < 0.01 and p-value<-2, recommended in DANPOS manual. The fuzziness of these nucleosomes may be increased or decreased from one condition to the other. We also used DNAPOS-generated wig files for visualization of Fuzziness on IGV genome browser.

**Mcd1 Pulldowns**

*MCD1-HA* and MCD1 untagged strains expressing CDC20 under the galactose promoter were grown at 25ºC in YEP medium containing 2% galactose to mid-logarithmic phase and arrested in metaphase by the addition of 2% glucose. 800 ODs of cells were harvested at 4ºC, washed once with ice-cold water and frozen at -80ºC. Cell pellets were mechanically broken in Lysis Buffer containing 150 mM KCl, 1mM MgCl2, 0.12mM Cacl2 50 mM Tris pH 7, 1 mM EDTA, 1% Triton 100-X, 5% glycerol and 1 mM PMSF supplemented with EDTA-free Complete Protease Inhibitor and Phosphatase Inhibitor Cocktail tablets (Roche). Protein extracts were treated with 600µg of RNase A (Qiagen), 900U of Benzonase (Millipore) and 600U of DnaseI (NEB) and immunoprecipitated using anti-HA Epitope Tag Protein Isolation kit (Miltenyi Biotec). After 1h incubation at 4ºC, anti-HA magnetic beads were washed with 750mM NaCl-Lysis Buffer and subjected to 2 consecutive rounds of elution in 50mM Tris-HCl pH6.8, 5mM EDTA, 1%SDS and 50mM DTT elution buffer.

**Protein identification and quantification by LC-MS/MS**

*Sample processing*-Protein samples eluted from anti-HA magnetic beads were buffer exchanged, reduced and alkylated using a FASP protocol. Briefly, samples were loaded onto 30 kDa centrifugal concentrators (Millipore, MRCF0R030) and buffer exchange was carried out by centrifugation on a bench top centrifuge (15min, 12,000g). Multiple buffer exchanges were performed sequentially with UA buffer (8M urea in 100mM Tris pH 8.5, 3x200□l), reduction with 10mM DTT in UA buffer (30min, 40°C) and alkylation with 50mM chloroacetamide in UA buffer (20min, 25°C). This was followed by buffer exchange into UA buffer (3x100□l) and 50mM ammonium bicarbonate (3x100□l). Digestion was carried out with mass spectrometry grade trypsin (Promega, V5280) using 1□g protease per digest (16h, 37°C). Tryptic peptides were collected by centrifugation into a fresh collection tube (10min, 12,000g) and washing of the concentrator with 0.5M sodium chloride (50□□l, 10min, 12,000g) for maximal recovery. Following acidification with 1% trifluoroacetic acid (TFA) to a final concentration of 0.2%, collected protein digests were desalted using Glygen C18 spin tips (Glygen Corp, TT2C18.96) and peptides eluted with 60% acetonitrile, 0.1% formic acid (FA). Eluents were then dried using vacuum centrifugation.

*Liquid chromatography-tandem mass spectrometry (LC-MS/MS) analysis-* Dried tryptic digests were redissolved in 0.1% TFA by shaking (1200rpm) for 30min and sonication on an ultrasonic water bath for 10min, followed by centrifugation (20,000g, 5°C) for 10min. LC-MS/MS analysis was carried out in technical duplicates and separation was performed using an Ultimate 3000 RSLC nano liquid chromatography system (Thermo Scientific) coupled to a coupled to a Q-Exactive mass spectrometer (Thermo Scientific) via an EASY spray source (Thermo Scientific). For LC-MS/MS analysis protein digest solutions were injected and loaded onto a trap column (Acclaim PepMap 100 C18, 100µm × 2cm) for desalting and concentration at 8µL/min in 2% acetonitrile, 0.1% TFA. Peptides were then eluted on-line to an analytical column (Acclaim Pepmap RSLC C18, 75µm × 50cm) at a flow rate of 250nL/min. Peptides were separated using a 120 minute gradient, 4-25% of buffer B for 90 minutes followed by 25-45% buffer B for another 30 minutes (composition of buffer B – 80% acetonitrile, 0.1% FA) and subsequent column conditioning and equilibration. Eluted peptides were analysed by the mass spectrometer operating in positive polarity using a data-dependent acquisition mode. Ions for fragmentation were determined from an initial MS1 survey scan at 70,000 resolution, followed by HCD (Higher Energy Collision Induced Dissociation) of the top 12 most abundant ions at 17,500 resolution. MS1 and MS2 scan AGC targets were set to 3e6 and 5e4 for maximum injection times of 50ms and 50ms respectively. A survey scan m/z range of 400 – 1800 was used, normalised collision energy set to 27%, charge exclusion enabled with unassigned and +1 charge states rejected and a minimal AGC target of 1e3.

*Raw data processing*- Data was processed using the MaxQuant software platform (v1.5.8.3), with database searches carried out by the in-built Andromeda search engine against the Uniprot S.cerevisiae database (version 20160815, number of entries: 6,729). A reverse decoy search approach was used at a 1% false discovery rate (FDR) for both peptide spectrum matches and protein groups. Search parameters included: maximum missed cleavages set to 2, fixed modification of cysteine carbamidomethylation and variable modifications of methionine oxidation, protein N-terminal acetylation and serine, threonine, tyrosine phosphorylation. Label-free quantification was enabled with an LFQ minimum ratio count of 2. ‘Match between runs’ function was used with match and alignment time limits of 1 and 20 minutes respectively.

#### Supplementary Figure legends

**Supplementary Figure 1.** Auxin-mediated degradation of Spt16 in $G_2$/M arrested cells. Cells carrying either *SMC1-6HA or SMC1-6HA SPT16-AID* (which allows auxin-mediated degradation of Spt16) were arrested in metaphase ($G_2$/M-nocodazole) and exposed to auxin to degrade Spt16. Nuclear and cellular morphology was used following auxin addition to confirm that the cell population remained arrested in metaphase (top graph). Western analysis for Spt16 confirms its degradation in *SPT16-AID* cells upon auxin addition (bottom).

**Supplementary Figure 2.** Auxin-mediated degradation of Sth1 in $G_2$/M arrested cells. Cells carrying either *SMC1-6HA or SMC1-6HA STH1-AID* (which allows auxin-mediated degradation of Sth16) were arrested in metaphase ($G_2$/M-nocodazole) and exposed to auxin to degrade Sth1. Western analysis for Sth1 confirms its degradation in *STH1-AID* cells upon auxin addition.

**Supplementary Figure 3.** FACT function in chromosome organisation.
The 16 yeast chromosomes are displayed on Hi-C maps (10 kb bins) obtained from $G_2$/M arrests in the presence of auxin of SMC1-6HA (wild-type, bottom left) and *SMC1-6HA SPT16-AID* (Spt16-aid) cells (top right). Black to white color scales reflect high to low contact frequencies, respectively ($log_2$). Inset display magnifications of chr7.
Analysis of chromosome segregation in the absence of FACT.

**Supplementary Figure 4.** FACT function in the establishment of cohesin-dependent structures in $G_1$.
The 16 yeast chromosomes are displayed on Hi-C maps (10 kb bins) obtained from $G_1$ arrests overexpressing *MCD1* from the *GAL1-10* promoter in the presence of auxin of *SMC1-6HA* (wild-type, bottom left) and *SMC1-6HA SPT16-AID* (Spt16-aid) cells (top right). Black to white color scales reflect high to low contact frequencies, respectively ($log_2$). Inset display magnifications of chr7.

**Supplementary Figure 5.** Analysis of chromosome segregation in the absence of FACT. *CDC15-AID* and *CDC15-AID SPT16-AID* cells were arrested in metaphase and exposed to auxin to degrade Cdc15 and Spt16 and then released from the metaphase arrest, by removing nocodazole, into telophase block caused by the absence of Cdc15 (Diagramme shows synchronisation - top left). The strains carried tetO/tetR-based chromosome tags inserted at right subtelomere region of chromosome IV that were used to score chromosome segregation (bottom graphs). Protein levels are shown (westerns- top right). Note that auxin addition leads to the degradation of Cdc15 and Spt16. Chromosome missegregation was observed in the absence of Spt16.

**Suppl. Figure 1**

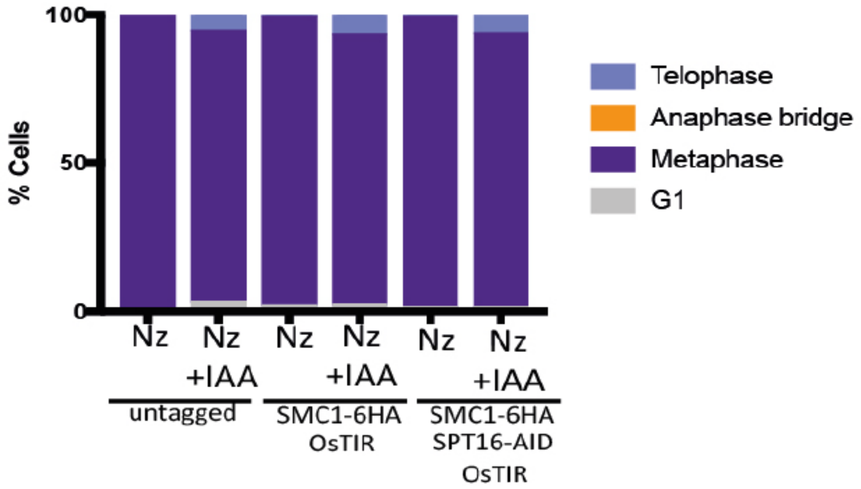


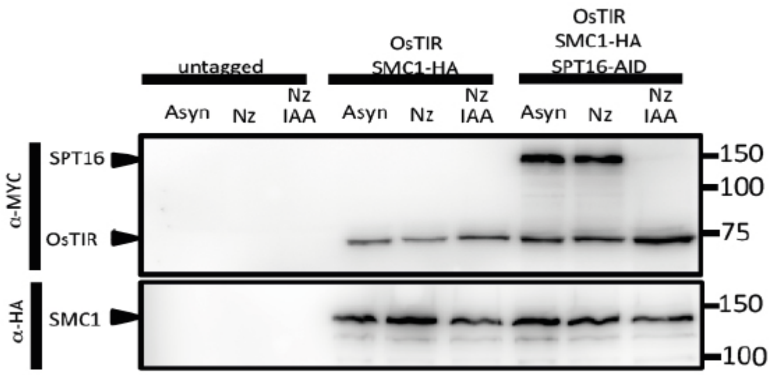

**Suppl. Figure 2**

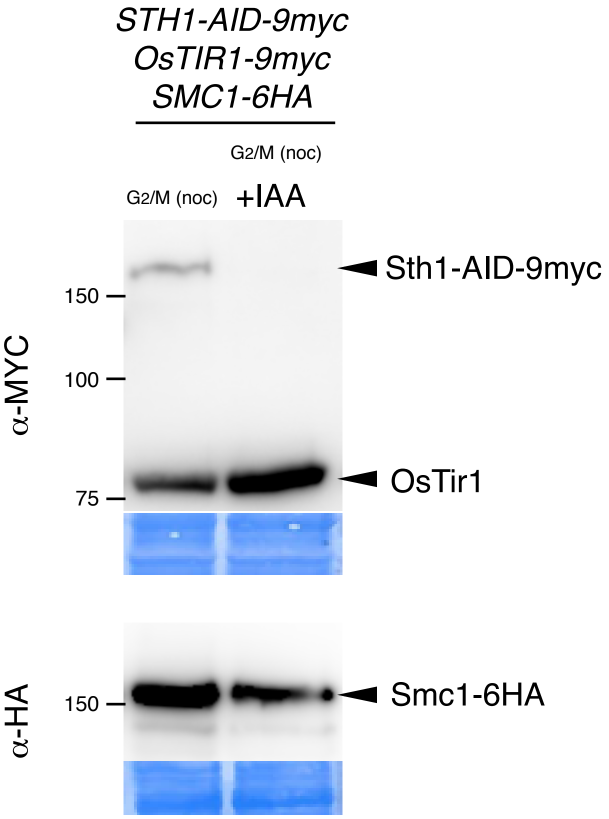

**Suppl. Figure 3**

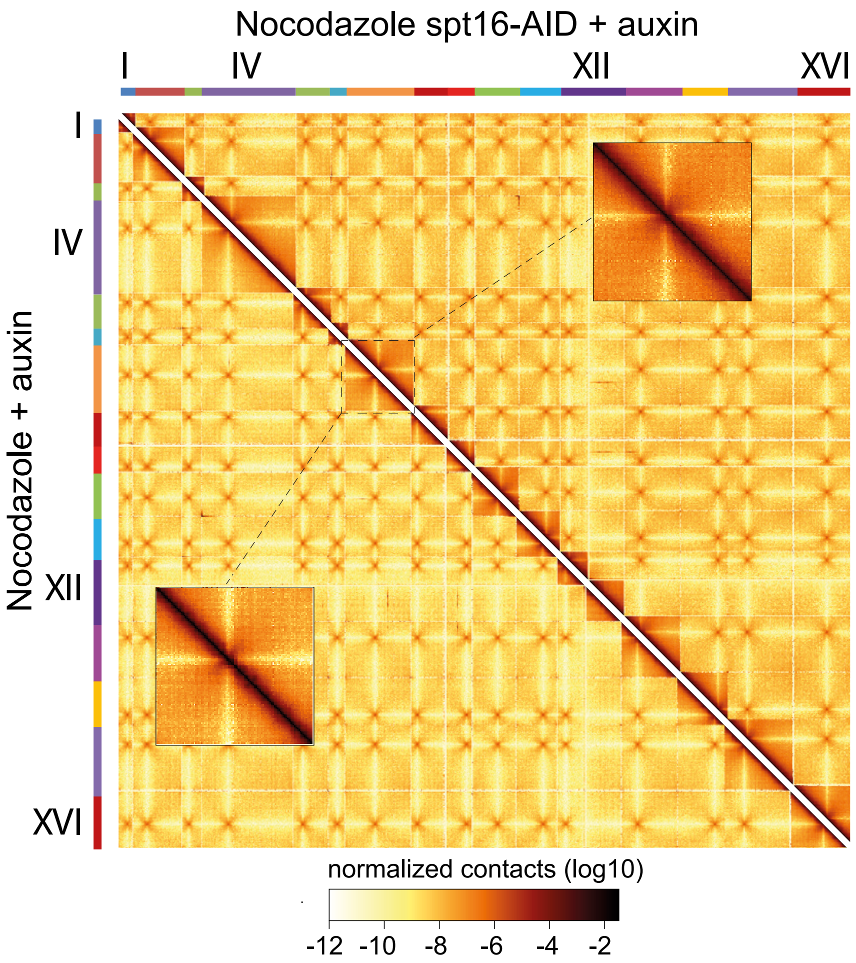

**Suppl. Figure 4**

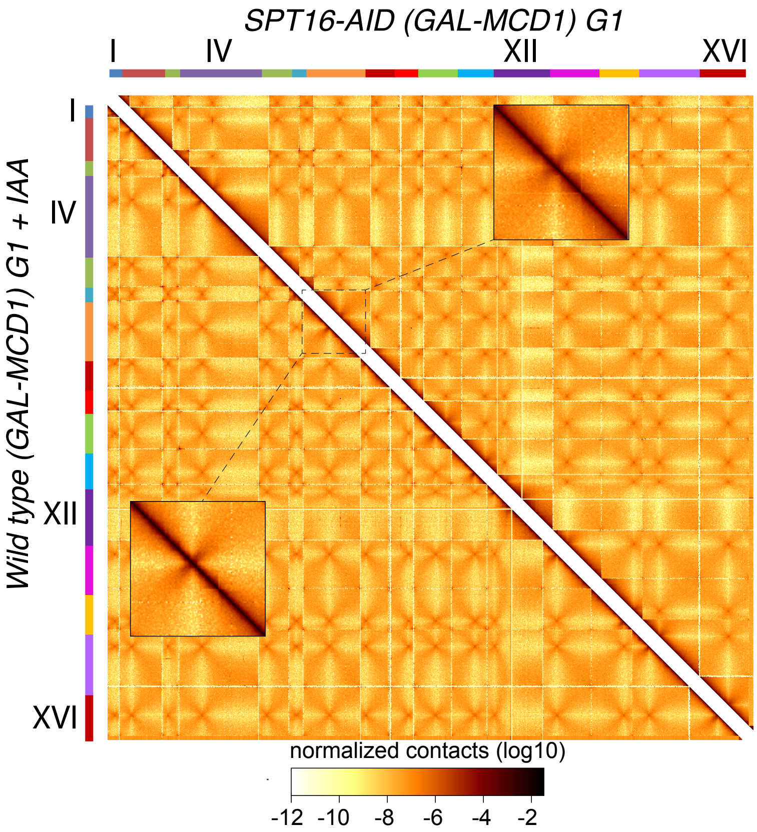

### Suppl. Figure 5

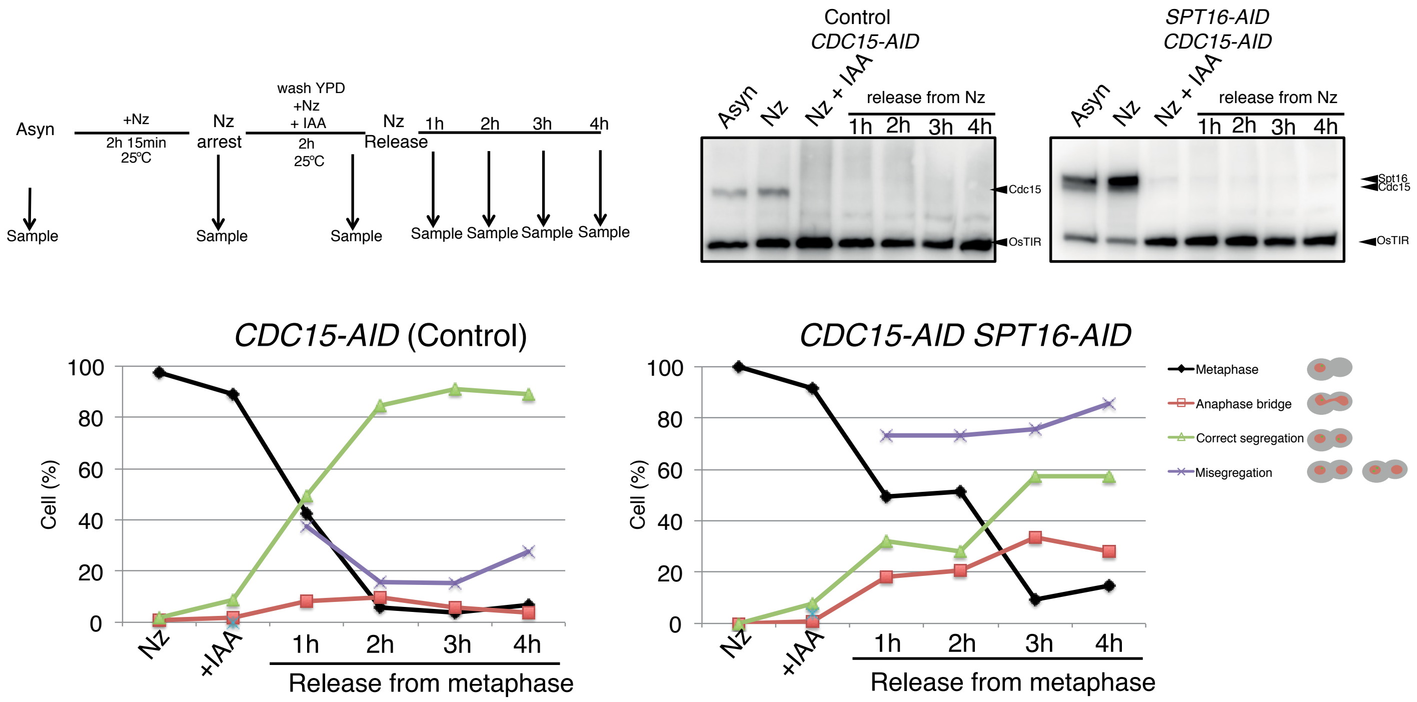
